# The miR-15a/16-1 and miR-15b/16-2 clusters regulate early B cell development by limiting IL-7 receptor expression

**DOI:** 10.1101/2022.03.18.484871

**Authors:** Katharina Hutter, Thomas Rülicke, Tamas G. Szabo, Lill Andersen, Andreas Villunger, Sebastian Herzog

## Abstract

Pleiotropic functions of miRNAs as transcriptional repressors have been reported for multiple biological processes. One prominent miRNA family is the miR-15 family, which is a well-established tumor-suppressor in B-cell chronic lymphocytic leukemia (CLL). The miR-15 family consists of three bicistronic clusters, miR-15a/16-1, miR-15b/16-2 and miR-497/195, all sharing the same seed sequence suggesting that loss one cluster can be functionally compensated by the remaining miR-15 family members. Thus, a combined deletion may be necessary to reveal its physiological function *in vivo*.

A combined knockout of the most prominent miR-15 clusters, miR-15a/16-1 and miR-15b/16-2 in the hematopoietic system reveals a novel role of the miR-15 family in early B cell development highlighted by an increase of the pro-B cell compartment. Mechanistically, this effect is mediated by enhanced IL-7 receptor expression, which we identified as direct miR-15 target gene. Notably, elevated IL-7 receptor levels were sufficient to trigger increased activation of the STAT5 and PI3K/AKT pathways. Moreover, derepression of directly targeted cell cycle regulators such as *Ccne1, Chek1* and *Wee1* further facilitates G-to-S transition.

Thus, by deregulating a target gene network of cell cycle and signaling mediators, loss of the miR-15 family establishes a pro-proliferative milieu manifesting in an enlarged pro-B cell pool.

## Introduction

Beyond the large network of transcription factors and chromatin regulators governing physiological as well as pathological processes, non-coding RNAs add another layer of gene regulation and thereby contribute to fine-tuning a large variety of biological pathways. MiRNAs as one well-studied group of non-coding RNAs exert their function by sequence-specific post-transcriptional repression of their target genes. This is mediated by binding to the 3’UTR of the respective target mRNA, resulting in inhibition of translation or mRNA decay (Bartel 2018). As miRNAs provide an important layer of gene regulation, it is not surprising that they are also involved in pathological conditions such as cancer (Di Leva et al. 2014). One prominent tumor-suppressive miRNA is the miR-15 family consisting of the three bicistronic clusters miR-15a/16-1, miR-15b/16-2 and miR-497/195 (Finnerty et al. 2010). The tumor-suppressive role of the miR-15a/16-1 cluster was first reported in B cell chronic lymphocytic leukemia (CLL), as 60 % of all patients exhibit a loss of the chromosomal region harboring this cluster (Calin 2002). This role of the miR-15 family in the CLL setting could be further recapitulated in mouse models, in which deletion of the miR-15a/16-1 or the miR-15b/16-2 cluster also promoted a CLL-like disease (Klein et al. 2010; Lovat et al. 2015). It is not surprising that both clusters are linked to CLL, as all clusters of the miR-15 family share the same seed sequence and thus regulate the same set of target genes (Bartel 2018). Noteworthy, whole-body deletion of miR-15a/16-1 and miR-15b/16-2 in mice did not result in accelerated CLL onset but rather induced acute myeloid leukemia (AML) development, indicating that distinct cell types have a different sensitivity for a reduction of miR-15 levels (Lovat et al. 2020). As the miR-497/195 cluster is only marginally expressed in immune cells, it is dispensable for immune cell development and function, while the other two clusters are highly expressed and several studies are beginning to elucidate their physiologic functions (Kuchen et al. 2010; Hutter et al. 2020). The miR-15a/16-1 cluster was shown to be important for natural killer (NK) cell development, as its NK cell-specific deletion partially blocked NK cell maturation in the spleen (Sullivan et al. 2015). Furthermore, a T cell specific combined deletion of the miR-15a/16-1 and miR-15b/16-2 cluster resulted in enhanced proliferation and accumulation of memory T cells upon lymphocytic choriomeningitis virus (LCMV) infection (Gagnon et al. 2019). Important functions of the miR-15 family have also been reported for early B cell development as an *in vitro* knockdown of the miR-15 family in pre-B cells resulted in impaired differentiation and enhanced proliferation (Lindner et al. 2017).

B cells develop in the bone marrow, where they undergo two sequential steps of immunoglobulin gene segment rearrangements, finally giving rise to a unique B cell receptor (BCR). Heavy chain diversity (D) to joining (J) rearrangement is initiated between the pre-pro and pro-B cell stage and is - after an initial proliferation wave - continued by variable (V) to DJ recombination in late pro-B cells (Schlissel 2003). Successful rearrangement results in an immunoglobulin heavy chain that pairs with the surrogate light chain, thus forming the pre-B cell receptor (pre-BCR) (Nishimoto et al. 1991). Expression of a fully functional pre-BCR induces several rounds of proliferation of large pre-B cells and subsequent light chain (VJ) recombination. This newly formed light chain can pair with the already existing heavy chain finally giving rise to a mature BCR, marking the immature B cell stage (Herzog et al. 2009). Immature B cells travel to the spleen where they undergo several additional maturation steps from transitional stage 1 (T1) to transitional stage 2 (T2) and stage 3 (T3), eventually ending up as mature marginal zone or follicular B cells (Chung et al. 2003; Pillai & Cariappa 2009). The two recombination events in early B cell development have to be tightly regulated to ensure proper separation of proliferation and somatic rearrangements, as errors in these processes compromise genomic stability and can result in malignant transformation and blood cancer (Küppers & Dalla-Favera 2001; Papaemmanuil et al. 2014; Zhang et al. 2011). One of the major regulatory signaling modules mediating proper segregation of the biphasic early B cell development is the IL-7 receptor and its downstream mediators STAT5 and the PI3K/AKT axis. IL-7 receptor signaling contributes to pro- and pre-B cell survival and proliferation and at the same time inhibits differentiation by downregulation of FOXO1 and repression of *Rag1* and *Rag2* genes. Thus, IL-7 receptor signaling has to be temporally turned off in order to allow pro- and pre-B cell differentiation, respectively (Clark et al. 2013). However, it is still not fully understood which molecular components establish this precise regulation throughout early B cell development.

Here we report a novel role of the miR-15 family in these processes. In particular, we demonstrate that combined loss of the miR-15a/16-1 and miR-15b/16-2 clusters in mice results in an increase of pro-B cells, indicating that one of the physiological functions of the miR-15 family is to restrict or limit expansion of B cell progenitors. Mechanistically, our data show that loss of the miR-15 clusters elevates the expression of the IL-7 receptor as a direct target gene, which is sufficient to result in enhanced activation and phosphorylation of IL-7 receptor downstream effectors such as AKT and STAT5. Additionally, deletion of the miR-15 family derepresses several cell cycle regulators such as *Wee1, Chek1, Ccnd3* and *Ccne1* in B cell progenitors, thereby facilitating G-to-S transition. Thus, enhanced growth factor receptor signaling combined with these transcriptional changes establish a pro-proliferative state, manifesting as an enlarged miR-15a/16-1 miR-15b/16-2 double knockout pro-B cell pool.

## Results

### Generation of conditional miR-15a/16-1 and miR-15b/16-2 double knockout mice

To study whether a complete loss of the miR-15 family members expressed in the hematopoietic compartment would affect physiologic immune cell development and function we generated a conditional miR-15a/16-1 miR-15b/16-2 knockout mouse model. We decided to neglect the miR-497/195 cluster in this setting, as it is only marginally expressed in immune cells and was found to be dispensable for the immune system in our previously described miR-497/195 knockout mouse model (Hutter et al. 2020). The conditional miR-15a/16-1 and miR-15b/16-2 knockout alleles were generated using CRISPR/Cas9-assisted editing to flank the endogenous clusters with loxP sites in embryonic stem (ES) cells (Figs. 1A and B). To delete the respective miRNA clusters in the hematopoietic system, mice derived from these ES cells were intercrossed and then bred with the Vav-Cre strain (Georgiades et al. 2002) to generate single miR-15a/16-1fl/fl (hereafter miR-15a^fl/fl Vav-Cre^) and miR-15b/16-2fl/fl (miR-15b^fl/fl Vav-Cre^) as well as miR-15a/16-1 miR-15b/16-2 (miR-15a/b^fl/fl Vav-Cre^) double knockout (DKO) mice.

**Figure 1.**
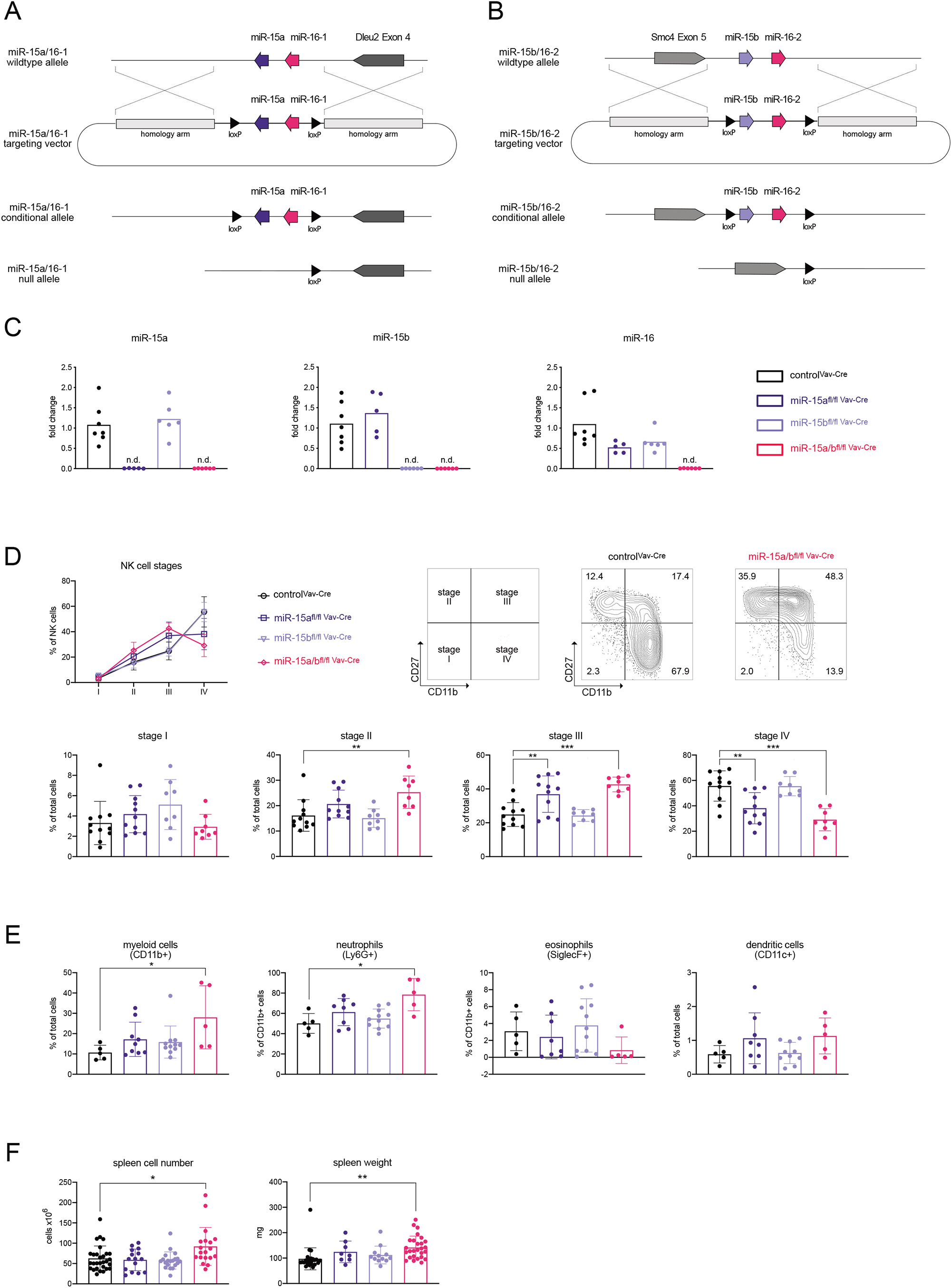
Generation of conditional miR-15a/16-1 and miR-15b/16-2 double knockout mice. (A and B) Targeting strategy for deletion of the miR-15a/-16-1 and miR-15b/16-2 cluster. The knock-in alleles were generated by CRISPR/Cas9-induced targeting of murine ES cells with a DNA template containing the miR-15a/16-1 or miR-15b/16-2 cluster flanked by loxP sites (black triangles) and two homology arms. Conditional miR-15a/16-1 or miR-15b/16-2 mice were crossbred to obtain miR-15a/16-1miR-15b/16-2 knockout mice. Cre-mediated deletion of the loxP-flanked DNA region creates the miR-15a/16-1 and miR-15b/16-2 null allele. (C) Quantitative PCR for the expression of mature miR-15a, miR-15b and miR-16 in B cells derived from miR-15a/b^+/+ Vav-Cre^, miR-15a^fl/fl Vav-Cre^, miR-15b^fl/fl Vav-Cre^ and miR-15a/b^fl/fl Vav-Cre^ mice. Each dot represents the data derived from one mouse. n.d. = not detected (D) NK cells defined as NK1.1+TCRb-were classified as stage I to stage IV based on their expression of CD27 versus CD11b (right side). Percentages of NK stage I to stage IV cells in the spleen of miR-15a/b^+/+ Vav-Cre^, miR-15a^fl/fl Vav-Cre^, miR-15b^fl/fl Vav-Cre^ and miR-15a/b^fl/fl Vav-Cre^ mice (left side and bar graphs). (E) Bar graphs show the percentages of indicated myeloid cells populations within total cells or CD11b+ myeloid cells. (F) Bar graphs indicate the total splenic cell numbers and spleen weight for the respective genotypes. Each dot represents the data derived from one mouse. Error bars depict the standard deviation of the mean. Each of the different miR-15 knockout groups were statistically compared to miR-15a/b^+/+ Vav-Cre^ mice by an unpaired two-tailed Student’s t-test. *P < 0.05; **P < 0.005; ***P < 0.0005.

To verify loss of the respective miRNAs in all genotypes compared to the control, we analyzed their expression in sorted B cells by quantitative PCR. As expected, we could observe a loss of miR-15a in miR-15a^fl/fl Vav-Cre^ and miR-15a/b^fl/fl Vav-Cre^ mice and a loss of miR-15b in miR-15b^fl/fl Vav-Cre^ and miR-15a/b^fl/fl Vav-Cre^ mice, respectively. Since the mature miR-16-1 and miR-16-2 are identical, a complete loss of miR-16 could only be detected in miR-15a/b^fl/fl Vav-Cre^ mice, whereas miR-15a^fl/fl Vav-Cre^ and miR-15b^fl/fl Vav-Cre^ mice showed a 50% reduction compared to miR-15a/b^+/+ Vav-Cre^ controls (Fig. 1C).

### Loss of miR-15 slightly enhances myeloid cell numbers and mildly affects the memory T cell compartment

To unravel a physiological function of the miR-15 family beyond its prominent role as tumor-suppressor in B-CLL, we first analyzed the non-B cell immune compartments in the conditional knockout mouse models under steady state conditions.

A previous study by Sullivan et. al described a partial block in NK cell development at the stage III to stage IV transition upon NK-cell specific deletion of the miR-15a/16-1 cluster (Sullivan et al. 2015). Validating the mouse model, our analysis recapitulated this finding, and interestingly this phenotype was even stronger in miR-15a/b^fl/fl Vav-Cre^ compared to the single knockout mice (Figs. 1D and S1A). Notable, while clearly contributing to the DKO phenotype, the miR-15b/16-2 cluster alone seems to be dispensable for NK cell maturation, as miR-15b^fl/fl Vav-Cre^ mice did not show any perturbations upon deletion of the cluster, pointing towards different roles of the individual miR-15 family clusters in this setting (Fig. 1D).

In contrast to NK cells, loss of either the miR-15a/16-1 or miR15b/16-2 cluster seems to mildly affect the myeloid compartment, as we observed only a slight, but not significant increase in myeloid cell frequencies. This elevation was further increased in DKO mice, revealing an expansion of neutrophils (Fig. 1E), probably resulting in a considerably higher spleen weight and splenic cellularity, as the total number of myeloid cells was significantly increased in miR-15a/b^fl/fl Vav-Cre^ mice (Figs. 1F and S1B). This is in accordance with a recent publication reporting the development of AML at an age of 5 months upon a combined whole body deletion of the miR-15a/16-1 and miR-15b/16-2 cluster (Lovat et al. 2020).

When analyzing T cell development in the thymus and the splenic T cell compartment, no major alterations upon loss of the miR-15 family were found under steady state conditions (Figs. 2A and B and S1C). The only deviation observed was a slight increase in CD8+ effector/memory T cell (T_EFF_) numbers and CD4+ central memory T cells (T_CM_) in miR-15a/b^fl/fl Vav-Cre^ mice, which supports a recent publication describing a role of the miR-15 family in restraining memory T cell differentiation, albeit in an infection model (Gagnon et al. 2019). Together, these data indicate a dose-dependent role of the miR-15a/16-1 and miR-15b/16-2 clusters in some, but not all non-B cell hematopoietic lineages, and overall validate the function of our mouse model. This, of course, raised the question about the effects of the combined deletion in B cells as the lineage mostly affected by pathological loss of one of the miR-15 family clusters.

**Figure 2.**
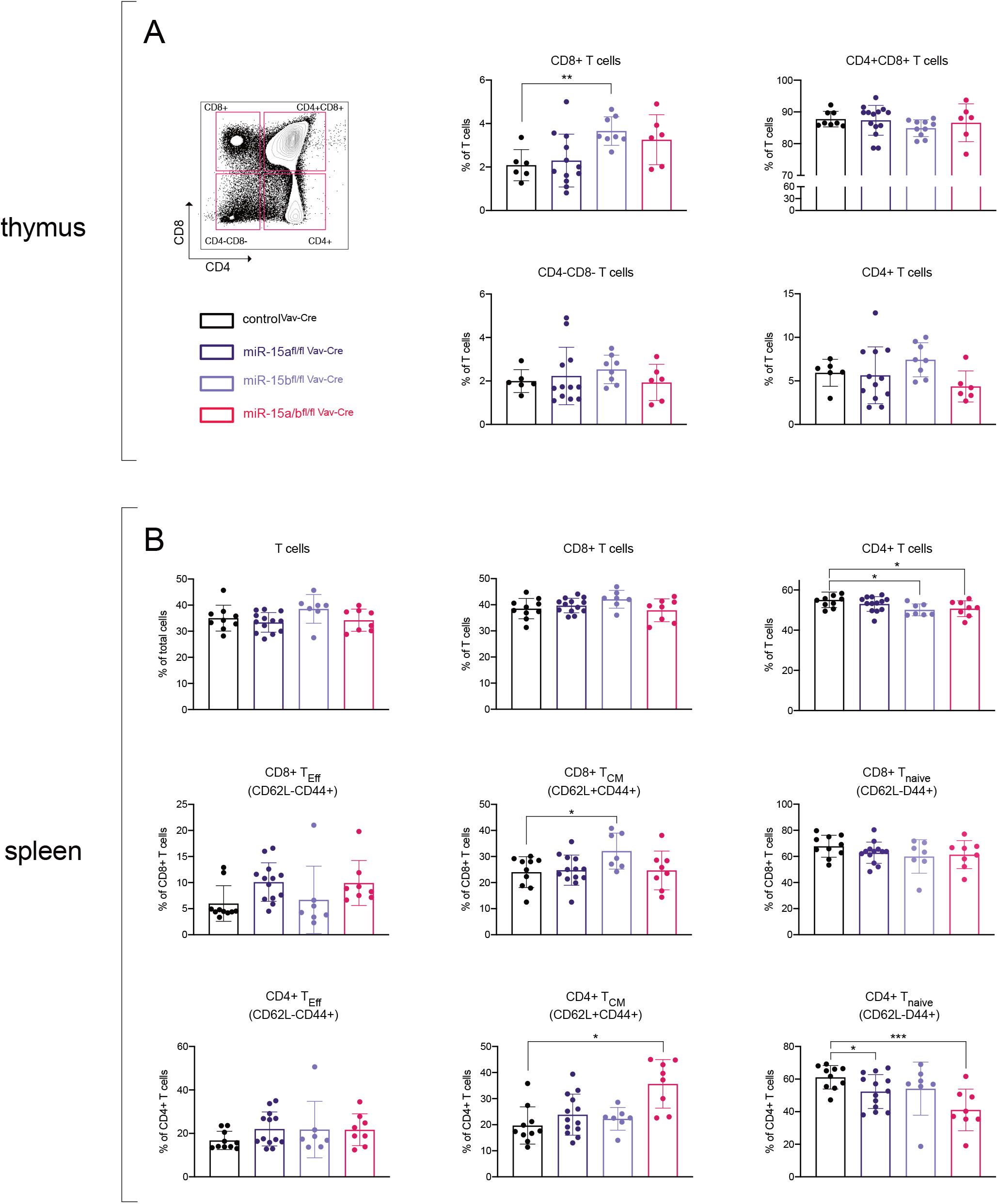
Minor changes in the T cell compartment upon loss of the miR-15 family. (A) Thymic T cells were defined as NK1.1-Gr-1-CD11b-B220-CD19-Ter119- and further divided into CD4+, CD8+, CD4+CD8+ and CD4-CD8-T cells (left side). Bar graphs show the percentages of respective thymic T cell populations within the T cell pool (right side). (B) Splenic CD8+ and CD4+ T cells were sub-gated into naïve, central memory and effector memory T cells according to CD62L and CD44 expression and respective percentages are depicted in bar graphs. Each dot represents the data derived from one mouse. Error bars depict the standard deviation of the mean. Each of the different miR-15 knockout groups were statistically compared to miR-15a/b^+/+ Vav-Cre^ mice by an unpaired two-tailed Student’s t-test. *P < 0.05; **P < 0.005; ***P < 0.0005.

### Loss of the miR-15a/16-1 and miR-15b/16-2 clusters impairs B cell development and maturation

Based on our own previous work pointing to an important role of the miR-15 family in early B cell development (Lindner et al. 2017), we initially analyzed the B cell composition of the bone marrow in miR-15 single and DKO mice (Fig. 3A). Overall, the total number of B cells in the bone marrow was reduced upon miR-15 loss, which is likely caused by the expansion of the myeloid cell pool (Figs. S2A). Within the B cell compartment, we could observe an increase in the pro-B cell subset for all three knockout genotypes (Fig. 3B), with the effect being most severe in the DKO mice. Correspondingly, the pre-B cell fractions were significantly increased in the single knockouts, but only by trend in the DKO, most likely due to the strong relative increase in pro-B cells (Fig. 3B). To test whether the observed effect of the miR-15 family on early B cell development is B cell intrinsic, we additionally generated a B cell-specific deletion of miR-15a/b, by crossing our miR-15a/b^fl/fl^ mice with the Mb1-Cre strain. As Mb1 encodes the BCR subunit Igα, it is expressed already at a very early pro-B cell stage, thus we could induce a combined deletion of the miR-15a/b clusters from the pro-B cell stage onwards (Hobeika et al. 2006). Similar to miR-15a/b^fl/fl Vav-Cre^ mice, B cell-specific deletion of miR-15a/b resulted in elevated pro-B cell frequencies in miR-15a/b^fl/fl Mb1-Cre^ mice, and in this context, pre-B cells were also significantly increased (Fig. S2B). This indicates that the early B cell phenotype is indeed B cell intrinsic. In this line, the total pro- and pre-B cell numbers also tended to be increased, albeit not significantly (Fig. S2C). To address whether the higher progenitor B cell percentages are derived from a general bias of hematopoiesis towards the lymphocyte lineage, we also examined the bone marrow regarding its composition of different hematopoietic lineage progenitors, i.e. common lymphoid progenitors (CLPs), granulocyte-monocyte progenitors (GMPs), megakaryocyte-erythroid progenitors (MEPs) and common myeloid progenitors (CMPs). However, these subsets did not show any alterations in miR-15a^fl/fl Vav-Cre^, miR-15b^fl/fl Vav-Cre^ and miR-15a/b^fl/fl Vav-Cre^ mice (Fig. S3A), pointing to a function of the miR-15 family only once committed to the B cell lineage.

**Figure 3.**
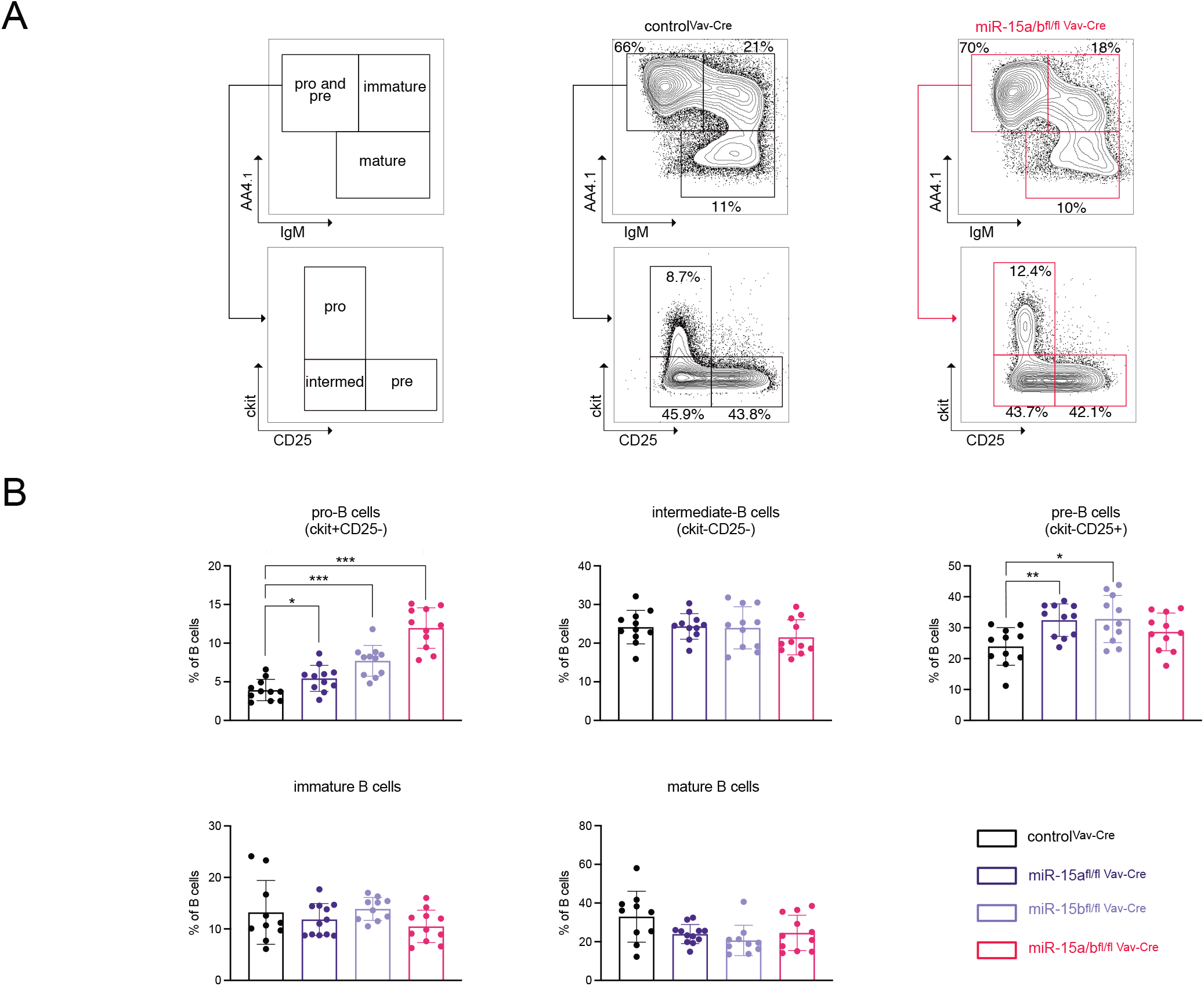
Loss of the miR-15a/16-1 and miR-15b/16-2 cluster impairs early B cell development. (A) Representative FACS plot illustrating the gating of pro-B and pre-B (AA4.1+IgM-), immature (AA4.1+IgM+) and mature B cells (AA4.1-) in the bone marrow. Pro-B and pre-B cells were further separated into pro-B (c-Kit+), intermediate-B (c-Kit-CD25-) and pre-B cells (CD25+). (B) Bar graphs show the percentage of pro-B, intermediate and pre-B cells (upper lane) and immature and mature B cells (lower lane) within the B220+CD19+ B-cell population in the bone marrow of miR-15a/b^+/+ Vav-Cre^, miR-15a^fl/fl Vav-Cre^, miR-15b^fl/fl Vav-Cre^ and miR-15a/b^fl/fl Vav-Cre^ mice. Each dot represents the data derived from one mouse. Error bars depict the standard deviation of the mean. Each of the different miR-15 knockout groups were statistically compared to miR-15a/b^+/+ Vav-Cre^ mice by an unpaired two-tailed Student’s t-test. *P < 0.05; **P < 0.005; ***P < 0.0005.

Having found this effect of miR-15 loss on early B cell development, we further examined later B cell developmental stages in the spleen (Fig. 4A). Beyond the early B cell stages, a reduction of miR-15 levels also affected mature B cell populations, as loss of the miR-15a/16-1 or miR-15b/16-2 or both clusters resulted in an increase in follicular B cells at the expense of marginal zone (MZ) B cell frequencies (Fig. 4B). These perturbations were not reflected in total cell numbers in miR-15a/b^fl/fl Vav-Cre^ mice (Fig. S4A), most likely due to a masking effect in response to the reported increase in the myeloid compartment. We therefore also analyzed the B cell-restricted miR-15a/b^fl/fl Mb1-Cre^ mouse model and could recapitulate the reduced percentages in MZ B cells, here accompanied by a reduction in total MZ B cell numbers (Figs. S4B and S4C). Thus, both miR-15a/16-1 and miR-15b/16-2 clusters are intrinsically required to efficiently establish the MZ B cell compartment, with a slightly stronger contribution of the former (Fig. 4B). Beyond the mature B cell compartment, a complete loss of miR-15 levels in miR-15a/b^fl/fl Mb1-Cre^ mice also led to a slightly skewed B cell maturation from transitional stage I towards stage II (Figs. 4B and S4B). However, these changes could only be observed as alterations within the immature B cell subset itself, as the total numbers of transitional B cells remained unchanged (Figs. 4B and S4C).

**Figure 4.**
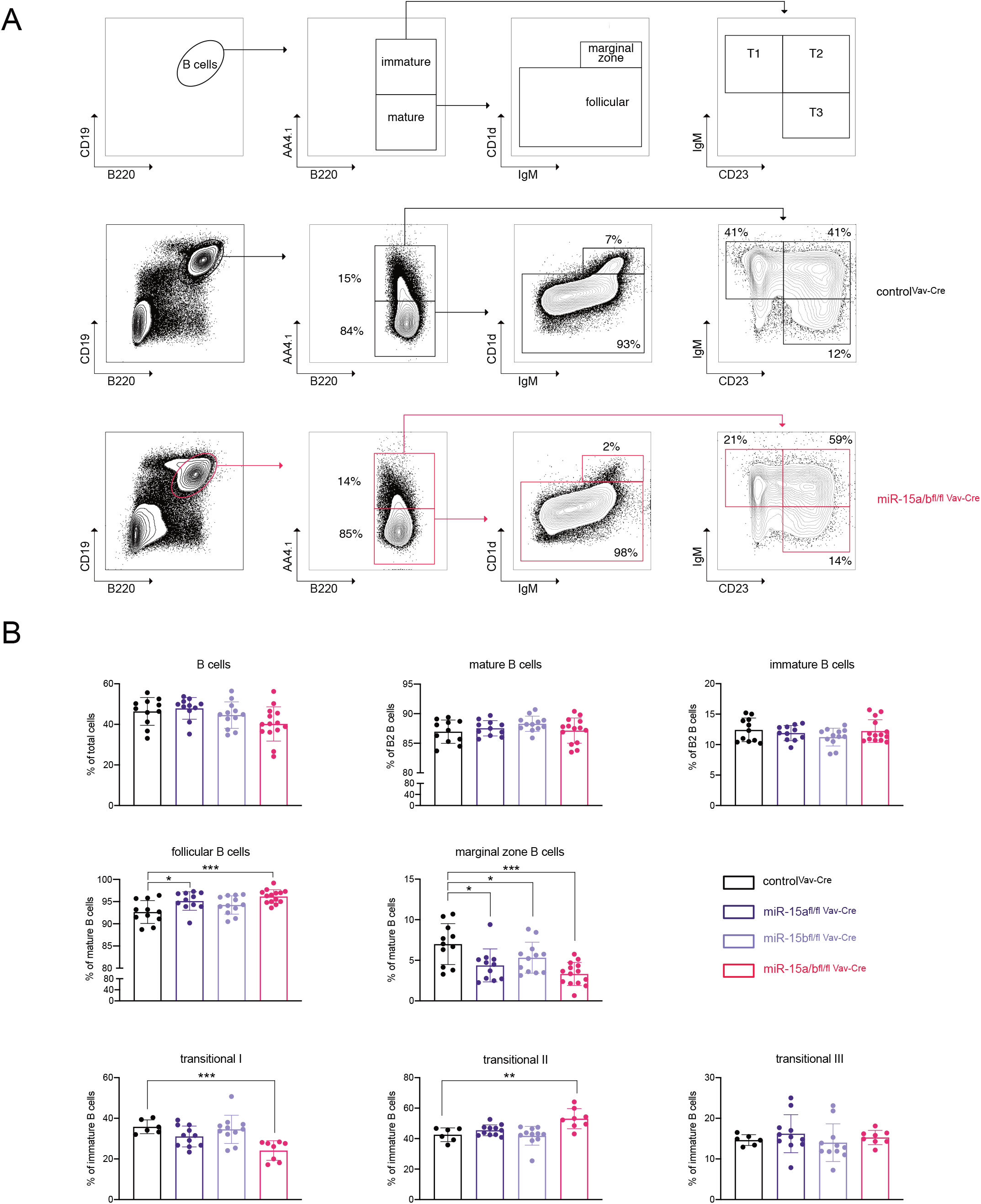
miR-15 levels control splenic B-cell development. (A) Gating strategy for the identification of different B-cell subsets in the spleen. CD19+B220+ B2 B cells were divided into mature (AA4.1-) and immature B cells (AA4.1+). The mature B cells were further split into CD1d+ marginal zone B cells and CD1d-follicular B cells. Immature B cells were gated for the different transitional phases T1 (IgM+CD23-), T2 (IgM+CD23+), and T3 (IgM-CD23+). (B) The bar graphs show the percentage of the indicated B cell populations among all splenic cells, mature B cells or the immature B cell pool in the spleens of miR-15a/b^+/+ Vav-Cre^, miR-15a^fl/fl Vav-Cre^, miR-15b^fl/fl Vav-Cre^ and miR-15a/b^fl/fl Vav-Cre^ mice. Each dot represents the data derived from one mouse. Error bars depict the standard deviation of the mean. Each of the different miR-15 knockout groups were statistically compared to miR-15a/b^+/+ Vav-Cre^ mice by an unpaired two-tailed Student’s t-test. *P < 0.05; *P < 0.005; ***P < 0.0005.

Taken together, these data clearly indicate that loss of the miR-15 family clusters affects early B cell development as well as later maturation stages in the spleen, pointing towards a crucial role in B cell physiology. However, with the early developmental phenotype being most prominent, we focused on characterizing the expansion of B cell precursors for the remainder of the study.

### Loss of the miR-15 family results in pro-B cell expansion *in vitro*

To investigate the role of the miR-15 family in early B cell development and to dissect the molecular mechanism underlying the miR-15-mediated regulation in more detail, we set up an *in vitro* pro-B cell culture system that allows the cultivation of primary B cell progenitors upon addition of IL-7, FLT3L and SCF (Muenchow et al. 2017)(Fig. 5A). Importantly, these progenitors still have the capability to further differentiate into distinct B cell developmental stages, resulting in a shift from intermediate (c-Kit^-^CD25^-^) towards pre-B cells (c-Kit^-^CD25^+^) upon IL-7 withdrawal (Fig. 5B). Exploiting this system, loss of the miR-15 family appeared to affect pre-B cell differentiation, albeit not statistically significant, with an altered intermediate to pre-B cell ratio in miR-15a/b^fl/fl Vav-Cre^ compared to control mice (Fig. 5C). Notably, steady state cell distributions were comparable, implying a direct effect on the differentiation process itself. As differentiation and proliferation appear to be mutually exclusive phases throughout early B cell development, we asked whether the reduced differentiation capacity and the elevated pro-B cell frequencies *in vivo* could be the consequence of enhanced expansion. To test this, we established an *in vitro* competition assay by labelling miR-15a/b^fl/fl Vav-Cre^ and control pro-B cells with different fluorescent markers and monitored the outgrowth of one or the other population over time upon co-culturing both genotypes. Interestingly, this resulted in a consistent expansion of miR-15a/b^fl/fl Vav-Cre^ cells over control cells, pointing towards increased survival, proliferation or both (Fig. 5D). To further assess this, we quantified EdU incorporation and analyzed apoptosis by AnnexinV staining. While the percentage of apoptotic cells was similar for control and miR-15a/b^fl/fl Vav-Cre^ cells, DNA synthesis as an indirect indicator for proliferation was significantly enhanced upon loss of the miR-15 family (Figs. 5E and F). Thus, our findings indicate that the miR-15a/16-1 and miR-15b/16-2 clusters limit pro-B cell proliferation and enable efficient differentiation under physiological conditions.

**Figure 5.**
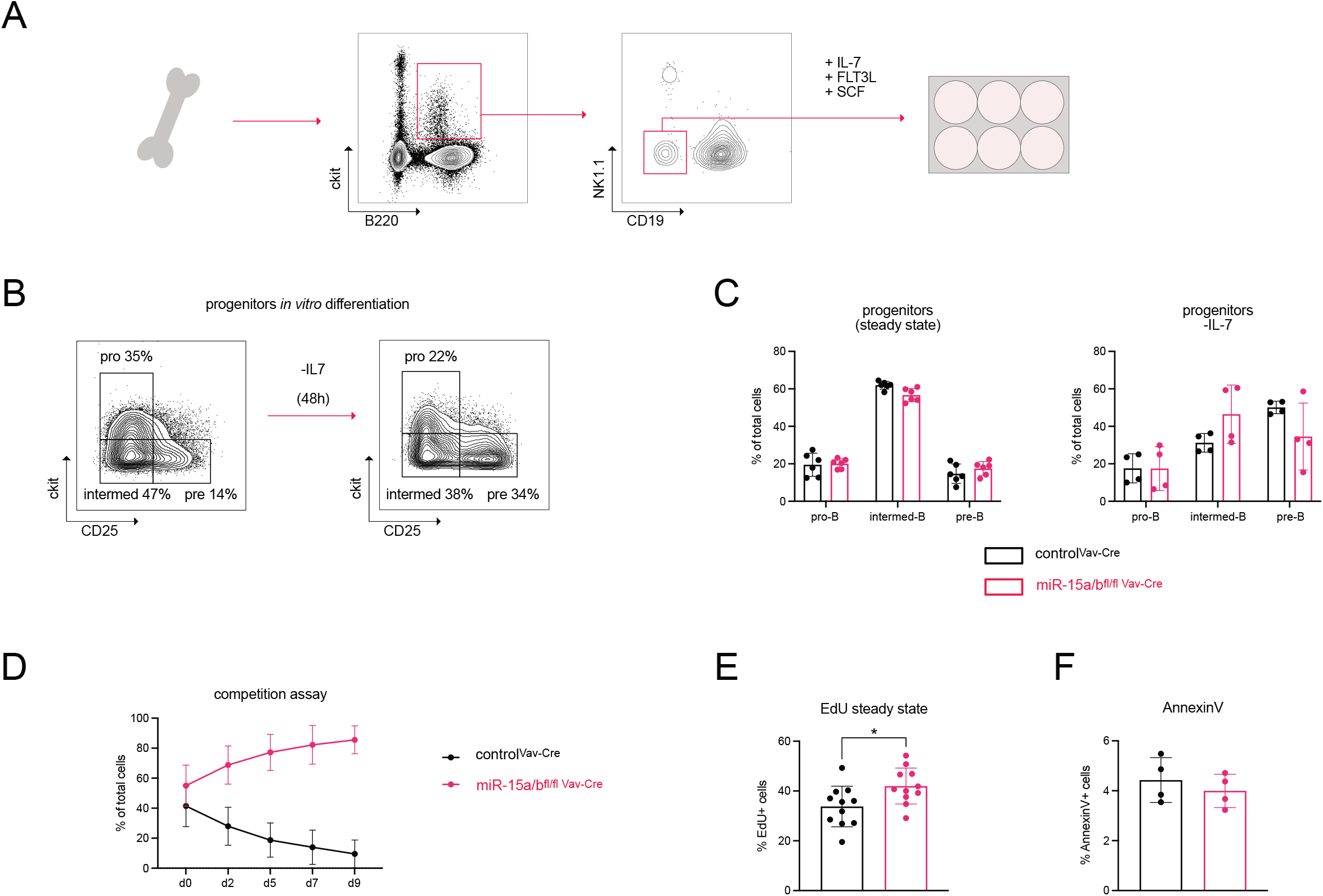
Loss of the miR-15 family results in pro-B cell expansion *in vitro*. (A) Schematic illustration of the sorting strategy for the B cell progenitor *in vitro* culture. B220+c-Kit+NK1.1-CD19-B cell progenitors were sorted from bone marrow of miR-15a/b^fl/fl Vav-Cre^ and miR-15a/b^+/+ Vav-Cre^ mice and cultured in the presence of IL-7, FLT3L and SCF. (B) Representative FACS plots showing the percentages of pro- (c-Kit+CD25-), intermediate (c-Kit-CD25-) and pre- (c-Kit-CD15+) B cells under basal and –IL-7 conditions. (C) Bar graphs summarize the quantification of pro-, intermediate and pre-B cell progenitors derived from miR-15a/b^fl/fl Vav-Cre^ mice and miR-15a/b^+/+ Vav-Cre^ mice. (D) Progenitors from miR-15a/b^fl/fl Vav-Cre^ mice and miR-15a/b^+/+ Vav-Cre^ mice were labelled by retroviral transduction for the expression of either dsRed or GFP, mixed at a specific ratio and the percentages of the respective groups were analyzed by flow cytometry. (E) The bar graph depicts the percentages of EdU+ progenitors derived from miR-15a/b^fl/fl Vav-Cre^ mice or miR-15a/b^+/+ Vav-Cre^ mice. 1 ug/ml EdU was added to the progenitors and EdU incorporation was detected after 4h by the ClickIt EdU method and flow cytometric quantification of EdU+ cells. (F) Quantification of the percentage of apoptotic progenitors under steady state conditions as analyzed by AnnexinV staining. Each dot represents the data derived from one experiment. Error bars depict the standard deviation of the mean. miR-15a/b^fl/fl Vav-Cre^ progenitors were statistically compared to miR-15a/b^+/+ Vav-Cre^ progenitors by an unpaired two-tailed Student’s t-test. *P < 0.05; **P < 0.005; ***P < 0.0005.

### Direct regulation of cell-cycle related genes by the miR-15 family

Having identified a clear function of the miR-15 family in restricting pro-B cell expansion, we next sought to find the target genes underlying this process. Therefore we performed RNA sequencing of sorted pro-B cells of control and miR-15a/b^fl/fl Vav-Cre^ mice and filtered all upregulated genes for conserved miR-15 binding sites using the well-established targetscan and miRDB algorithms (Agarwal et al. 2015; Chen & Wang 2019; Liu & Wang 2019). This resulted in the identification of 65 putative miR-15 target genes (Figs. 6A and B), which were strongly enriched for processes involved in cell cycle progression (Fig. 6C). In consequence, we further narrowed down the list of promising putative upregulated target genes to the ones that are known as direct or indirect cell cycle regulators. To examine, whether these genes could be indeed direct targets of the miR-15 family, we performed 3’UTR reporter assays in a pre-B cell line (Fig. 6D). Among the 3’UTRs that showed GFP repression upon miR-15a/16-1 overexpression were some prominent cell cycle regulating genes such as *Ccne1, Wee1, Cdc25* and *Chek1*, some of which have already been described as miR-15 family-regulated targets (Lindner et al. 2017; Mei et al. 2015; Sueur et al. 2020). To validate these findings, the upregulation of a selected set of genes in response to miR-15 knockout was confirmed by qPCR analysis (Fig. 6E). Together, these data indicate that the miR-15 family limits early B cell development at least in part by regulating critical cell cycle mediators.

**Figure 6.**
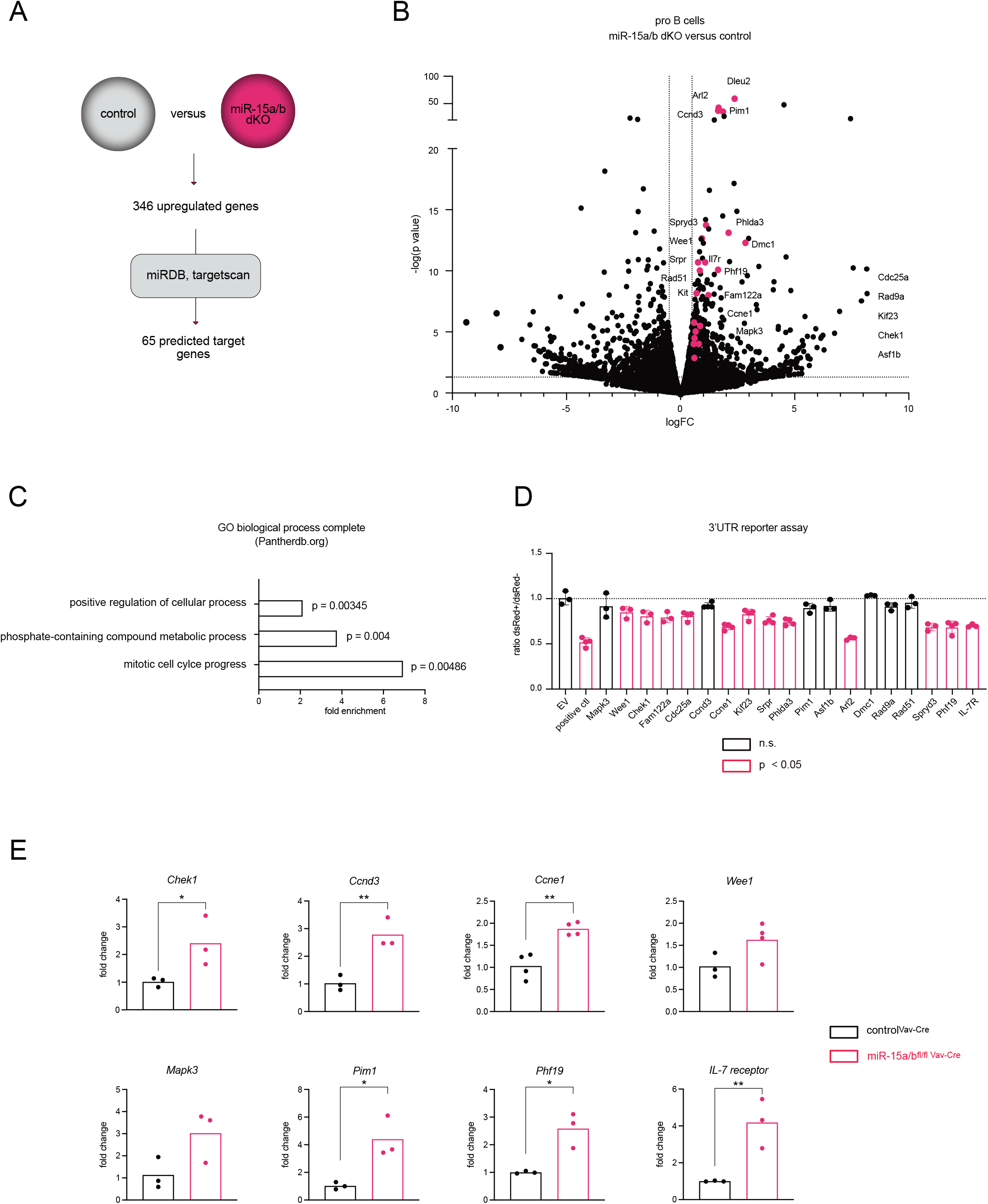
The miR-15 family directly regulates IL-7 receptor expression and cell-cycle related genes. (A) Schematic illustration of the identification of putative miR-15 target genes identified by RNA sequencing. Upregulated genes in pro-B cells of miR-15a/b^fl/fl Vav-Cre^ mice compared to miR-15a/b^+/+ Vav-Cre^ mice were filtered for predicted miR-15a, miR-15b and miR-16 binding sites in their 3′-UTRs using miRDB and targetscan. (B) Volcano plot depicting the up- and downregulated genes analyzed by RNA sequencing. The pink dots mark genes that contain a putative miR-15 binding site and have been selected for further 3’UTR reporter assays. (C) Gene Ontology analysis summarizing the strongest enriched processes in miR-15a/b^fl/fl Vav-Cre^ compared to miR-15a/b^+/+ Vav-Cre^ pro-B cells. (D) Pre-B cells were selected for the expression of a reporter construct encoding GFP with the 3′-UTR of the indicated target genes. Cells expressing GFP without a 3′-UTR served as negative control, whereas cells with a 3′-UTR containing two perfect miR-16 target sites were defined as positive control. The bars highlighted in pink depict 3’UTRs that led to significant GFP repression upon miR-15 overexpression. Error bars depict the standard deviation of the mean. The different groups were compared to the no-3′-UTR-negative control by a one-way ANOVA followed by Bonferroni testing (E) Quantitative PCR for the expression of selected genes in pro-B cells of miR-15a/b^fl/fl Vav-Cre^ mice compared to miR-15a/b^+/+ Vav-Cre^ mice. Each dot represents the data derived from one mouse. The deltaCT values of miR-15a/b^fl/fl Vav-Cre^ mice were statistically compared to miR-15a/b^+/+ Vav-Cre^ mice by an unpaired two-tailed Student’s t-test. *P < 0.05; **P < 0.005; ***P < 0.0005.

### The IL-7 receptor is a direct target of the miR-15 family

Beside the targets with a direct role in cell cycle regulation, we also retrieved the IL-7 receptor as one of the upregulated genes in miR-15a/b^fl/fl Vav-Cre^ pro-B cells (Figs. 6B, D and E). Notably, IL-7 receptor signaling is well-established as a key regulator of early B cell survival, proliferation and differentiation, with both pro- and pre-B cells expressing high levels of the receptor on their surface (McLean & Mandal 2020). Accordingly, CRISPR-Cas9-mediated knockout of the IL-7 receptor in progenitor B cells resulted in rapid depletion of the cells (Fig. S5) underlining the critical role of this pathway. The IL-7 receptor 3’UTR contains two miR-15 binding sites that are conserved across species (Fig. 7A). To address the function of both sites individually we mutated either the first or the second or both binding sites and performed 3’UTR reporter assays as mentioned above. In addition to the miR-15 overexpression setting, we decided to induce a knockdown of the miR-15 family by transducing pre-B reporter cells with a miR-15 sponge construct containing multiple miR-15 binding sites and thereby sequestering specific miRNA/RISC complexes. Loss of either of the two binding sites already de-repressed GFP reporter expression upon miR-15 overexpression, which was further enhanced upon mutating both binding sites. Conversely, knockdown of the miR-15 family resulted in increased GFP expression, which was reduced upon mutating either of the two binding sites. Furthermore, this GFP upregulation was completely abolished by a combined loss of both binding sites (Fig. 7B), clearly indicating that the IL-7 receptor mRNA is directly targeted through both miR-15 binding sites. To investigate whether the increased mRNA levels also translate onto the protein level, thus being functionally relevant, we analyzed IL-7 receptor surface expression of different B cell developmental stages in our miR-15a/b^fl/fl Vav-Cre^ mice as well as on progenitors *in vitro*. Indeed, loss of the miR-15 family resulted in enhanced IL-7 receptor surface expression in all analyzed settings (Figs. 7C and D). Since STAT5 is an important downstream element of IL-7 receptor signaling, mediating proliferation and survival of early B cells (Clark et al. 2013), we analyzed a potential impact of elevated IL-7 receptor expression on its phosphorylation. While direct flow cytometric measurements of activating pSTAT5 levels in both genotypes turned out to be difficult, probably due to low basal levels under steady state conditions, pSTAT5 induction was significantly enhanced in miR-15 DKO progenitors compared to the control when cells were starved and then stimulated with a defined amount of IL-7 (Fig. 7E). Extending our analysis to PI3K/AKT signaling, another pathway downstream of the IL-7 receptor that mediates survival and maintenance of pro- and pre-B cells (Clark et al. 2013), we furthermore found increased activating pAKT protein levels in progenitors derived from miR-15 DKO mice compared to controls (Figs. 7F and G). This points towards an increased activation of both the STAT5 as well as the PI3K/AKT axes downstream of the IL-7 receptor even under unaltered IL-7 conditions, likely contributing to the observed B cell progenitor expansion *in vivo*. Notably, this phenotype may be further strengthened by increased surface expression of c-KIT upon loss of miR-15 family expression, an additional receptor with a role in pro-B cell survival and expansion (Fig. 7H). However, as the *c-Kit* 3’UTR does not contain miR-15 binding sites, this upregulation is most likely not a direct effect, but is mediated through another, yet to be identified target gene.

**Figure 7.**
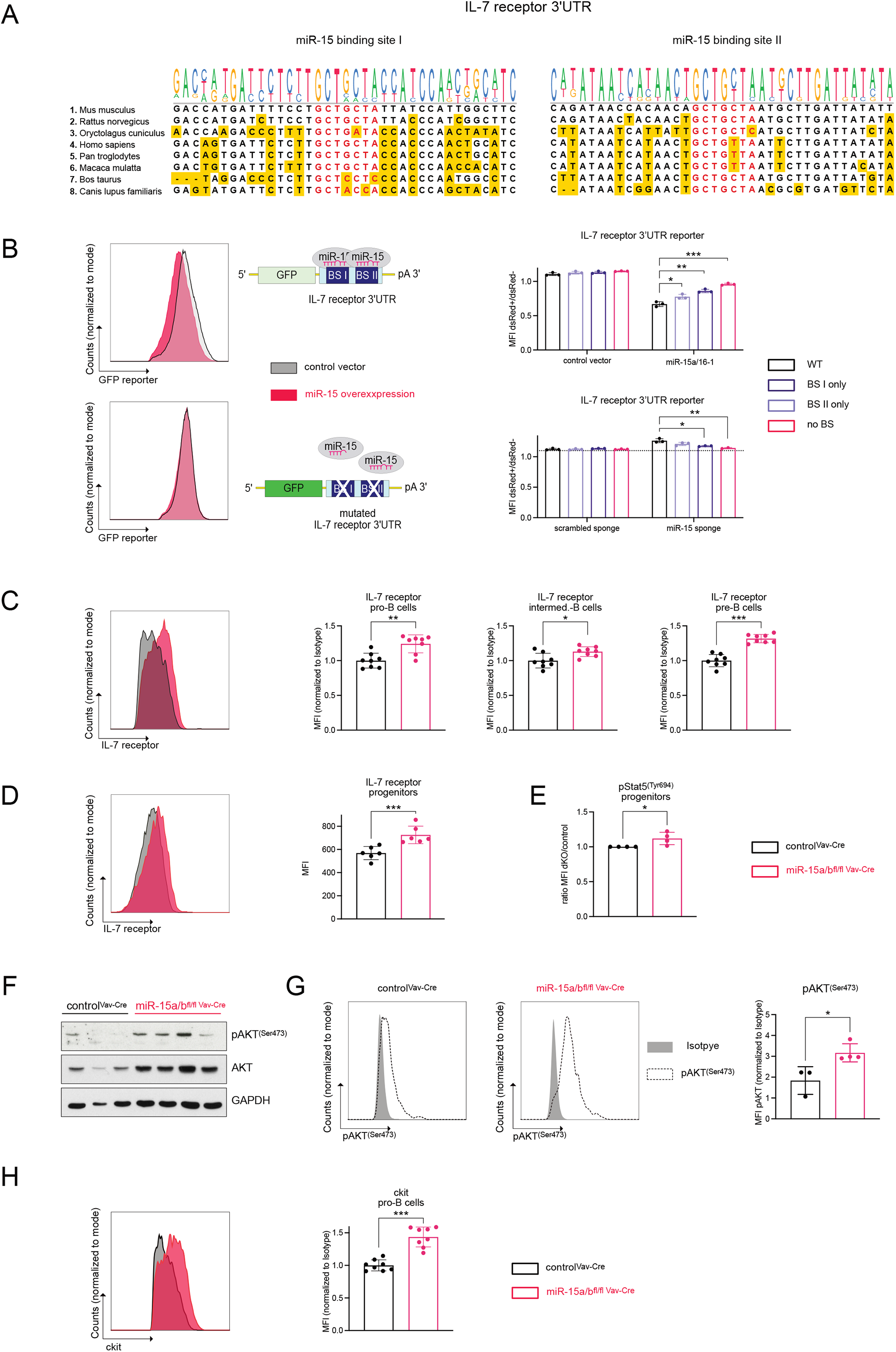
The IL-7 receptor is a direct target of the miR-15 family. (A) Schematic illustration of the two IL-7 receptor miR-15 binding sites in different species. The yellow highlighted nucleotides differ from the murine locus. (B) Representative histograms depicting GFP expression of the wild type and mutated (BS1+II) IL-7 receptor 3’UTR constructs upon miR-15 overexpression (left side). Pre-B cells expressing GFP linked to the IL-7 receptor 3’UTR or to the IL-7 receptor 3’UTR having mutations in the miR-15 BS I (BS II only) or BS II (BS I only) or both binding sites (no BS). Cells were transfected with either a dsRed construct, dsRed-miR-15a/16-1, dsRed-scrambled sponge or dsRed-miR-15 sponge. The height of the bars indicates the ratio of GFP MFI comparing transduced and non-transduced cells as measured by flow cytometry (right side). The miR-15 overexpression or miR-15 sponge samples were statistically compared to empty vector or scrambled sponge controls by an unpaired two-tailed Student’s t-test. (C and D) IL-7 receptor expression of pro- (c-Kit+CD25-), intermediate- (c-Kit-CD25-) and pre- (c-Kit-CD25+) B cells from control or miR-15a/b^fl/fl Vav-Cre^ mice as well as *in vitro* cultured progenitors, as analyzed by flow cytometry. The MFI values were normalized to the respective isotype control. (E) pSTAT5^(Tyr694)^ levels upon IL-7 starvation for 1 hour followed by addition of 0.1 ng/ul IL-7 for 30 minutes. The MFIs of anti-pSTAT5^(Tyr694)^ stained cells were normalized to the respective Isotypes. (F) Western blot analysis of control and miR-15a/b^fl/fl Vav-Cre^ progenitors depicting pAKT^(Ser473)^, AKT and GAPDH protein levels. (G) Representative histograms of flow cytometric analysis of intracellular pAKT^(Ser473)^ levels in control and miR-15a/b^fl/fl Vav-Cre^ progenitors. The bar graph summarizes the pAKT^(Ser473)^ MFI normalized to the respective Isotype. (H) Representative histogram (left) and summarizing bar graph (right) depicting the flow cytometric analysis of c-Kit surface expression normalized to the respective Isotype in miR-15a/b^fl/fl Vav-Cre^ or miR-15a/b^+/+ Vav-Cre^ pro-B cells. Each dot represents the data derived from one experiment. miR-15a/b^fl/fl Vav-Cre^ progenitors were statistically compared to miR-15a/b^+/+ Vav-Cre^ progenitors by an unpaired two-tailed Student’s t-test. *P < 0.05; **P < 0.005; ***P < 0.0005.

## Discussion

Given the high levels of miR-15a, miR-15b and miR-16 in various immune cell subsets, but the comparably low expression of miR-497/195 herein, we sought to study the physiological role of the miR-15 family by combined loss of the miR-15a/16-1 and miR-15b/16-2 clusters (Kuchen et al. 2010; Spierings et al. 2011). Exploiting this mouse model, we were able to identify immune cell compartments that are particularly sensitive to a small decrease in miR-15 levels, as it occurs in either miR15a/16-1 or miR-15b/16-2 single knockout mice, as well as cells that tolerate deletion of one cluster but are severely affected by the loss of the whole miR-15 family. With respect to the former, we could confirm the previous finding that NK cells are exceptionally dependent on high miR-15 levels, since deletion of the miR-15a/16-1 cluster alone already resulted in a maturation block (Sullivan et al. 2015). Interestingly, loss of the miR-15b/16-2 cluster did not alter NK cell maturation, however, a combined loss of both clusters further aggravated the developmental block at the stage III to IV transition. While we cannot exclude that this is due to a slightly distinct set of miR-15a target genes in contrast to miR-15b, a more likely explanation is that miR-15a/16-1 is simply expressed to higher extent in this subpopulation. If so, it is plausible that loss of miR-15b/16-2 does not reduce miR-15 levels below the critical threshold to provoke a phenotype, unlike the loss of the higher expressed miR-15a/16-1. However, deletion of both clusters in such a setting completely abolishes miR-15 family expression, and consequently strengthens the developmental delay.

In the T cell compartment, we could detect slight alterations with respect to CD8+ T cell percentages in the thymus, CD4+ T cell frequencies in the spleen and differences in central memory versus naïve T cell subsets. Minor changes in the T cell compartment of miR-15a/b^fl/fl Vav-Cre^ mice are not unexpected and in accordance with a recent study reporting a restrained T cell memory formation upon LCMV infection in combined miR-15a/16-1 and miR-15b/16-2 knockout mice (Gagnon et al. 2019).

With respect to its tumor suppressive role, the miR-15 family has been connected to several hematologic malignancies including CLL, in which more than 60% of patients display a genomic deletion comprising the miR-15a/16-1 cluster. Accordingly, loss of either miR-15a/16-1 or miR-15b/16-2 in mouse models has been linked to B-CLL development upon aging (Klein et al. 2010; Lovat et al. 2015). Surprisingly, however, combined whole body deletion of miR-15a/16-1 and miR-15b/16-2 has been reported to result in AML development at an age of 5 months (Lovat et al. 2020). Supporting this finding, our data also revealed increased numbers of myeloid cells in spleen and bone marrow already in young miR-15a/b DKO mice. Mice lacking either cluster were only slightly affected, suggesting that myeloid cells can cope with deletion of one cluster, but not with the loss of the whole miR-15 family. For the spleen, we could observe an increase in neutrophils which is probably responsible for the expanded myeloid compartment. However, additional studies will be needed to further investigate the role of the miR-15 family in myeloid cells, and to identify the target genes that mediate its tumor suppressive role in AML. One interesting candidate in this context is

CDC25A, which has been reported to be regulated by miR-16 in an AML subtype (FLT3-ITD AML) and serves as a critical factor for AML development (Sueur et al. 2020). In accordance with the already described role of miR-16 in *Cdc25a* repression, our data confirm *Cdc25a* as direct miR-15 target gene in the B cell setting. It will therefore be interesting to follow-up on this axis in myeloid cells and in the context of AML in the miR-15 DKO mouse model.

In the B cell compartment, our data reveal a role of the miR-15 family in marginal zone (MZ) B cell generation and/or maintenance, as a deletion of one or both clusters reduced the overall numbers of MZ B cells in a dose-dependent manner. As the main determinant of MZ versus follicular B cell fate decision appears to be BCR signaling strength, with weaker signals favoring the development of MZ B cells, one possibility of miR-15 function in this setting could be the dampening of BCR downstream signaling pathways (Pillai & Cariappa 2009). Alternatively, mutations involving the BAFFR and the canonical NF-κB pathway, the NOTCH pathway and integrin or chemokine signaling have all been associated with loss of the MZ compartment (Shulga-Morskaya et al. 2004; Cariappa et al. 2000; Saito et al. 2003; Lu & Cyster 2002). It is therefore possible that loss of miR-15-mediated regulation interferes with any of these processes, however, these hypotheses need to be further addressed in future studies.

Besides the MZ B cell reduction, the main phenotype observed in the B cell compartment was a strong increase in progenitor B cells, indicating a restrictive function of the miR-15 family under physiological conditions. A role of the miR-15 family in early B cell development has already been shown *in vitro*, where a functional knockdown of the miR-15 family in a pre-B cell line facilitated proliferation and counteracted pre-B to immature B cell differentiation. Mechanistically, the miR-15 family was reported to exert its functions via direct and indirect targeting of the cell cycle regulators *cyclin E1* and *cyclin D3*, respectively (Lindner et al. 2017). In consequence, cells lacking miR-15 family function failed to exit the cell cycle, which appeared critical to induce the transcriptional program required for differentiation. In the *in vivo* experiments presented here, B cell development was most severely altered at the pro-B cell stage in our miR-15a/b DKO knockout mice, however, pre-B cell percentages were elevated as well, in the single and in the double DKO mice. A partial block at both stages, pro- and pre-B cells, is not unexpected, as the pro- to pre-B transition requires a similar set of genes as the subsequent pre-B to immature B cell differentiation (Clark et al. 2013). Supporting this, we also found *cyclin E1* and *cyclin D3* to be strongly upregulated upon miR-15 deletion in pro-B cells, together with other cell-cycle regulators such as *Wee1, Chek1* and *Cdc25a*, most of which have also been described in pre-B cells lacking miR-15 function. Notably, such a general dysregulation of the cell-cycle program has been also reported in B-CLL, non-small lung cancer and breast cancer samples with low miR-15 expression, further underlining the importance of the miR-15 family in establishing a proper cell-cycle control (Bandi et al. 2009; Luo et al. 2013; Klein et al. 2010). Notably, as both pro-B and pre-B cells have to stop proliferating and exit the cell-cycle in order to start their heavy and light chain rearrangement, respectively, it is likely that the dysregulated cell-cycle progression alone counteracts further differentiation.

An additional key target of the miR-15 family in this context appears to be the IL-7 receptor gene, which we found strongly upregulated in the pro-B cell setting and confirmed in gain-of-function and loss-of-function 3’-UTR assays. In consequence, surface expression of the IL-7 receptor was clearly elevated in pro-as well as in pre-B cells. All of these progenitor B stages are well known to be highly responsive to IL-7, and IL-7 receptor signaling is critical for survival and cell expansion throughout early B cell development (Clark et al. 2013; Peschon et al. 1994; Grabstein et al. 1993). Notably, data in pre-B cells indicate that the miR-15 family not only controls IL-7 receptor expression, but is also regulated by IL-7 receptor signaling itself, pointing towards a crucial role of this axis in early B cell development (Lindner et al. 2017).

In terms of signaling downstream of the IL-7 receptor, our data indicate a clear activation of the JAK1/3/STAT5A as well as of the PI3K/AKT pathways. both of which known to contribute to pro- and pre-B cell survival and proliferation (Clark et al. 2013). In particular, stimulated miR-15 DKO-derived progenitors exhibited enhanced STAT5 phospho-Tyr694 levels *in vitro*, most likely reflecting elevated IL-7 receptor expression. On the other hand, Ser473 phosphorylation of AKT, which is considered as a marker for pathway activation, was strongly increased in miR-15a/b DKO progenitors even under steady state conditions. Our data therefore add another miRNA family to the growing list of PI3K regulators, which is not surprising in the light of this pathway’s key function in early B cell development (Ochiai et al. 2012). In addition to the miR-15 family, another important group of miRNAs essential for B cell differentiation beyond the pro-B cells stage is the miR-17-92 cluster, which targets *Pten* and thereby regulates PI3K activity (Benhamou et al. 2018). Likewise, the miR-26 family, another miRNA family with a role in early B cell development, directly represses *Pten* in pre-B cells (unpublished data). Hence, miRNAs modulate PI3K activation on different levels, thereby controlling the developmental progression of B cell progenitors.

Last, our data also provide evidence for an increased expression of *c-Kit* upon deletion of the miR-15 clusters, and it is tempting to speculate that this enhanced expression also contributes to the pro-proliferative phenotype. Indeed, several *in vitro* studies indicate a stimulatory role of signaling via c-KIT in pro-B cells (McNiece et al. 1991; Rolink et al. 1991; Yasunaga et al. 1995), however, B cells lacking *c-Kit* expression have been shown to develop normally (Takeda et al. 1997). While this points to a non-essential role of *c-Kit* in lymphopoiesis, it does not exclude a positive impact once upregulated, in particular in the context of *c-Kit* being considered as a proto-oncogene in several cancer entities (Liang et al. 2013). Thus, it remains to be determined whether the increased *c-Kit* expression is a bystander effect without any functional relevance, or whether it in fact aggravates the progenitor B cell phenotype induced by loss of miR-15 family members.

Taken together, our data confirm and build on previous findings regarding the role of the miR-15 family in NK cell maturation, memory T cell maintenance and myeloid expansion, and establish a novel function in early B cell development *in vivo*. In correspondence with its described tumor-suppressive function, physiological levels of miR-15 family members restrict or limit expansion of progenitor B cells at several stages, thereby ensuring proper transitions between phases of proliferation and differentiation. As it has been described in other cellular systems such as e.g. T cells (Gagnon et al. 2019), miR-15 expression appears to exert this role by modulating an extensive target gene network instead of regulating one specific master gene. Thus, we propose that the increase in the pro- and pre-B cell compartments upon miR-15 family loss is mediated via multiple pathways, including aberrant cell-cycle regulation as well as elevated growth factor receptor expression, which in synergy orchestrate the enhanced proliferation and compromised differentiation.

## Material and Methods

### Ethics statement

Experimental procedures with animals were discussed and approved by the institutional ethics and animal welfare committees of the University of Veterinary Medicine Vienna and the Medical University of Innsbruck in accordance with good scientific practice guidelines and national legislation (license numbers: BMBWF-68.205/0023-II/3b/2014 and BMBWF-66.011/0021-V/3b/2019).

### Animals

The conditional miR-15a/16-1 and miR-15b/16-2 alleles were generated by CRISPR/Cas9-facilitated homologous recombination in murine ES cells, as already described for targeting of the miR-497/195 cluster (Hutter et al. 2020). In short, KH2 ES cells ((Premsrirut et al. 2011), C57BL/6 × 129S4/Sv/Jae background, kindly provided by J. Zuber, IMP, Vienna) were electroporated (Nucleofector Kit; Lonza, Switzerland) with two Cas9/sgRNA vectors encoding GFP as a marker and the targeting DNA template containing the miRNA cluster flanked by loxP sites. After 36 h, ES cells were sorted for GFP^hi^ cells and plated at a low density on feeder cells. Individual ES cell clones were screened by PCR, sequenced, and then used for injection in C57BL/6NRj blastocysts. High percentage chimeras were bred with C57BL/6NRj females to confirm germline transmission and then further backcrossed to generate a congenic strain. Cre-mediated recombination in hematopoietic stem cells was induced by mating with C57BL/6N.Cg-Tg(Vav-Cre) mice (Georgiades et al. 2002) or C57BL/6N.Cg-Tg(Mb1-Cre) mice to mediate B-cell specific deletion (Hobeika et al. 2006).

Animals were kept specific pathogen-free according to FELASA recommendations ((Mähler et al. 2014)) under controlled environmental conditions (temperature 22 °C ± 1 °C, relative humidity of 40– 60%), a 12:12-h light/dark cycle, in a facility for laboratory rodents. Food (regular mouse diet (Ssniff, Germany) and water were provided *ad libitum*. Mice were maintained in small groups in individually ventilated cages lined with wood shavings as bedding and enriched with nesting material. If not stated otherwise, mice were analyzed at an age of 10–12 weeks. For all experiments, male and female mice were used in comparable numbers.

### Preparation of single-cell suspension

Single-cell suspensions for flow cytometry were obtained by pulpifying spleens through a 70-µm filter. For bone marrow cell suspensions, femurs and tibiae were isolated, ground, and filtered through a 70-µm filter. Lysis of erythrocytes for spleen and blood samples was performed by incubating the cells for 3–5 min in 1 mL lysis buffer (155 mm NH_4_Cl, 10 mm KHCO_3_, 0.1 mm EDTA; pH 7.5). Cells were resuspended and washed in FACS buffer (PBS with 1% FCS). Cells were counted using a Neubauer counting chamber.

### Flow cytometry

Single-cell suspensions were stained in 96-well plates with 30 µL of the antibody cocktails for 20 min at 4 °C. Nonspecific antibody binding was blocked by pre-incubating the cells with anti-CD16/31 antibodies in 30 µL FACS buffer for 10 min at 4 °C. All centrifugation steps were performed with 530 g for 2 min. For the antibody cocktails, the following antibodies were used: anti-B220-BV510 (BioLegend, 103247, 1 : 300), anti-CD19-BV605 (BioLegend; 115540, 1 : 300), anti-AA4.1-PE/Cy7 (BioLegend; 136507, 1 : 300), anti-AA4.1-APC (BioLegend; 136510, 1 : 300), anti-CD25-PE (BioLegend; 102007, 1 : 300), anti-c-Kit-APC (BioLegend; 135108, 1 : 300), anti-CD1d-PE (Thermo Fisher Scientific; 12-0011-82, 1 : 400), anti-CD23-PE/Cy7 (BioLegend; 101614, 1 : 300), anti-IgM-FITC (BioLegend; 406506, 1 : 300), anti-IgD-PerCpCy5.5 (BioLegend; 405710, 1 : 300), anti-TCR-FITC (eBioscience; 11-5961-85, 1 : 300), anti-CD4-APC-Cy7 (BD Biosciences; 552051, 1 : 300), anti-CD8-BV421 (BioLegend; 100753, 1 : 300), anti-CD62L-PE (BD Biosciences; 553151, 1 : 400), anti-CD44-BV510 (BD Biosciences; 563114, 1 : 300), anti-NK1.1-APC (BioLegend; 108709, 1 : 300), anti-gdTCR-PE (BioLegend; 118108, 1 : 400), anti-streptavidin-BV605 (BioLegend; 405229, 1 : 400), anti-CD27-FITC (BioLegend; 124207, 1 : 300), anti-CD43-PE (BD Biosciences; 7297616, 1 : 300), anti-NK1.1-PeCy7 (BioLegend; 108714, 1 : 300), anti-C11b-APC-Cy7 (BioLegend; 101226, 1 : 300), anti-TCRbeta-PerCPCy5.5 (BioLegend; 109228, 1 : 300), and anti-B220-FITC (BioLegend; 103206, 1 : 300), anti-CD127-APC (BioLegend; 135020, 1 : 300), anti-IgG2a,k Isotype control (BioLegend; 400511, 1 : 300), anti-CD34-FITC (Miltenyi; 130-105-831, 1 : 300), anti-Sca1-PE (BioLegend; 108107, 1 : 300), anti-APC-Cy7-Streptavidin (BioLegend; 405208, 1 : 400). The lineage cocktail to exclude non-T cells in the thymus combined anti-B220-bio (eBioscience; 13-0452-75, 1 : 100), anti-CD19-bio (BioLegend; 115504, 1 : 100), anti-Gr-1-bio (eBioscience; 13-5931-75, 1 : 100), anti-NK1.1-bio (BioLegend; 108704, 1 : 100), anti-CD11b-bio (eBioscience; 13-0112-75, 1 : 100), and anti-Ter119-bio (eBioscience; 13-5921-75, 1 : 100) antibodies. The lineage cocktail to exclude mature immune cell populations for the hematopoietic progenitors analysis combined anti-B220-bio, anti-Ter119-bio, anti-CD11b-bio, anti-Gr-1-bio, anti-CD3-bio (eBioscience; 13-0031-75, 1 : 100).

Apoptotic cells were stained using Topro3 (1:10000) for exclusion of dead cells and AnnexinV-FITC (1:10000) prior to flow cytometric analysis.

Flow cytometric analysis was performed on a FACS LSR Fortessa instrument (BD Biosciences), and sorting was performed on a FACS Aria III (BD Biosciences) cell sorter. Data were analyzed using FlowJo software (BD Life Science, USA).

### Intracellular Staining (pAKT, pSTAT5)

After the surface staining, cells were washed in 1% BSA/PBS and fixed in 4% PFA/PBS for 15min at RT. After one washing step (1% BSA/PBS) cells were permeabilized in PermBuffer (saponin-based permeabilization and wash buffer; ThermoFisher) for 15min and subsequently stained with pAKT Ser473 (CellSignaling; 4060, 1 : 200), pSTAT5 Tyr694 (CellSignaling; 4322, 1: 200) or IgG Isotype control (CellSignaling; 3900; corresponding concentration to pAKT or pSTAT5) for 60min at RT. Cells were washed and incubated with the secondary antibody anti-rabbit IgG-Alexa647 (CellSignaling; 4414, 1 : 500) for 30min at RT.

### Quantitative real-time PCR

RNA of sorted cells was isolated using the Quick-RNA Microprep Kit according to manufacturer’s instructions (Zymo; R1050). For analysis of miRNA genes RNA was reverse-transcribed with the miRCURY LNA RT Kit (Qiagen, Hilden, Germany; 339340) followed by SYBR Green qPCR (miRCURY LNA SYBR Green PCR Kit; Qiagen; 339356). For RNA sequencing target validation, RNA was reverse-transcribed using the iScript cDNA synthesis Kit (Bio-rad 1708891) and the Luna Universal qPCR Master Mix (NEB, M3003). Final quantification was performed using the ΔΔCT method.

### RNA Sequencing

RNA sequencing was performed as a service by Novogene (Cambridge, UK). In short, total RNA was isolated using the RNeasy Micro Kit according to manufacturer’s instructions (Qiagen; 74004) and amplified and reverse transcribed into cDNA using the SMART-seq protocol (Takara), followed by a quality control and library preparation. Paired end sequencing (reads of 150 nt) was performed on a NovaSeq 6000 (Illumina). For data analysis, single end reads were trimmed using Trimmomatic (Bolger et al. 2014), aligned using STAR aligner (Dobin et al. 2013) and counted with FeatureCounts (Liao et al. 2014). Differential gene expression was analyzed using EdgeR (Robinson et al. 2010; McCarthy et al. 2012). (Data analysis conducted by Tamas Szabo).

### Transfections and viral transductions

Prior to transductions, retro- or lentiviral supernatants were produced in HEK293T cells. Per sample, 400 ng of DNA for the plasmid of interest, 150 ng of either pSPAX2 (for lentiviral plasmids) or HIT60 (for retroviral plasmids) and 150 ng pVSVg were diluted in 20 μl IMDM medium (Sigma-Aldrich) and mixed with 20 μl IMDM medium containing 1.2 μl of 1 mg/mL polyethylenimine (PEI, Polysciences). Following a 20 min incubation step at room temperature, the DNA:PEI mix was pipetted into a 24-well tissue culture plate and 230 μl of HEK293T at a density of 6 × 10^5^ cells per ml were plated on top. After 30 – 40 hours viral supernatants were harvested, mixed with polybrene (16 μg/mL final concentration) and used for spin-infection of cells in 1.5 ml reaction tubes or 96-well plates (400 g at 37°C for 90 min).

### 3’UTR reporter assays

UTRs of interest were amplified by PCR, purified by agarose gel isolation (Monarch DNA Gel Extraction Kit, NEB (T1020L)) and ligated into LMP-destGFP (Lindner et al. 2017) 3′ of the destabilized GFP cDNA via *XhoI* and *ClaI*. The pre-B cell line WK3 was transduced with the respective viral supernatants and cells were selected for puromycin resistance by adding puromycin at a concentration of 1 μg/ml for 3–4 days. In experimental settings, cells were always cultured in the absence of puromycin.

### Cell lines

#### Pre-B cell line for 3’UTR reporter assays

The pre-B cell line WK3 was derived by extended culture of total bone marrow of SLP-65-deficient mice in IL-7-supplemented medium (Lindner et al. 2017). Pre-B cells were cultured in IMDM (Sigma) containing 7.5% FCS (Biochrom Superior), 100 U/ml penicillin, 100 U/ml streptomycin (PAN), and 50 μM 2-ME. Pre-B cell medium was supplemented with IL-7 by addition of the supernatant of IL-7 secreting J558L cells.

#### B cell progenitors

Bone marrow of miR-15a/b^+/+Vav-Cre^ or miR-15a/b^fl/flVav-Cre^ mice was sorted for c-Kit+B220+CD19-NK1.1-B cell progenitors. Progenitors were cultured in IMDM containing 7.5% FCS (Biochrom Superior), 100 U/ml penicillin, 100 U/ml streptomycin (PAN), 50 μM 2-ME, 100ng/ml SCF, IL-7 supernatant and FLT3L supernatant). To induce differentiation of progenitors *in vitro*, cells were cultured in the absence of IL-7 for 48 hours.

For CRISPR/Cas9 induced targeting of the IL-7 receptor, Cas9-GFP expressing progenitors derived from B6(C)-Gt(ROSA)^26Sorem1.1(CAG-cas9*,-EGFP)Rsky^/J mice (Chu et al. 2016) were transduced with constructs encoding sgRNAs targeting the IL-7 receptor and dsRed as a marker.

#### EdU assay

(Click-iT(tm) EdU Alexa Fluor(tm) 647 Flow Cytometry Assay Kit; Invitrogen; C10419) For *in vitro* proliferation assays, 10ug/ml EdU was added to the medium and cells were harvested after 4 hours of incubation. After surface staining, cells were fixed in 100ul 4% PFA for 15 minutes at room temperature in a 96-well plate. All following steps were performed as according to manufacturer’s instruction, however the amounts of reagents used was downscaled to a 96-well format.

#### Western Blot

For Western blot analysis, cells were lysed in ice-cold RIPA buffer (50 mM Tris–HCl, pH 7.4, 1% NP-40, 0.25% sodium deoxycholate, 150 mM NaCl, 1 mM EDTA (pH 8), protease inhibitor cocktail Sigma, 10 mM Na3VO4, 10 mM NaF), mixed 1:1 with reducing sample buffer (62.5 mM Tris pH 6.8, 2% SDS, 10% Glycerol, 100 mM DTT), boiled for 5 min at 95°C, and load onto a SDS–PAGE gel. Western blotting was performed using PVDF membranes.

Proteins were detected using anti-pAKT Ser473 (CellSignaling; 4060), anti-AKT (Cell Signaling; C67E7), and anti-actin antibodies (CellSignaling; 13E5). Bound antibodies were visualized with HRP-labeled secondary antibodies and the ECL system (Advansta) on a light-sensitive film (Amersham, GE).

#### Statistical analysis

Values in figures depict mean ± standard deviation (SD), and each mouse or experiment is represented as a dot. The two experimental groups were statistically analyzed by unpaired two-tailed Student’s t-tests using prism 9 software (GraphPad). P-values < 0.05 were considered as statistically significant, and graphs were labeled according to following scheme: ***P < 0.005, **P < 0.005, and *P < 0.05.

---

**Figure S1.**
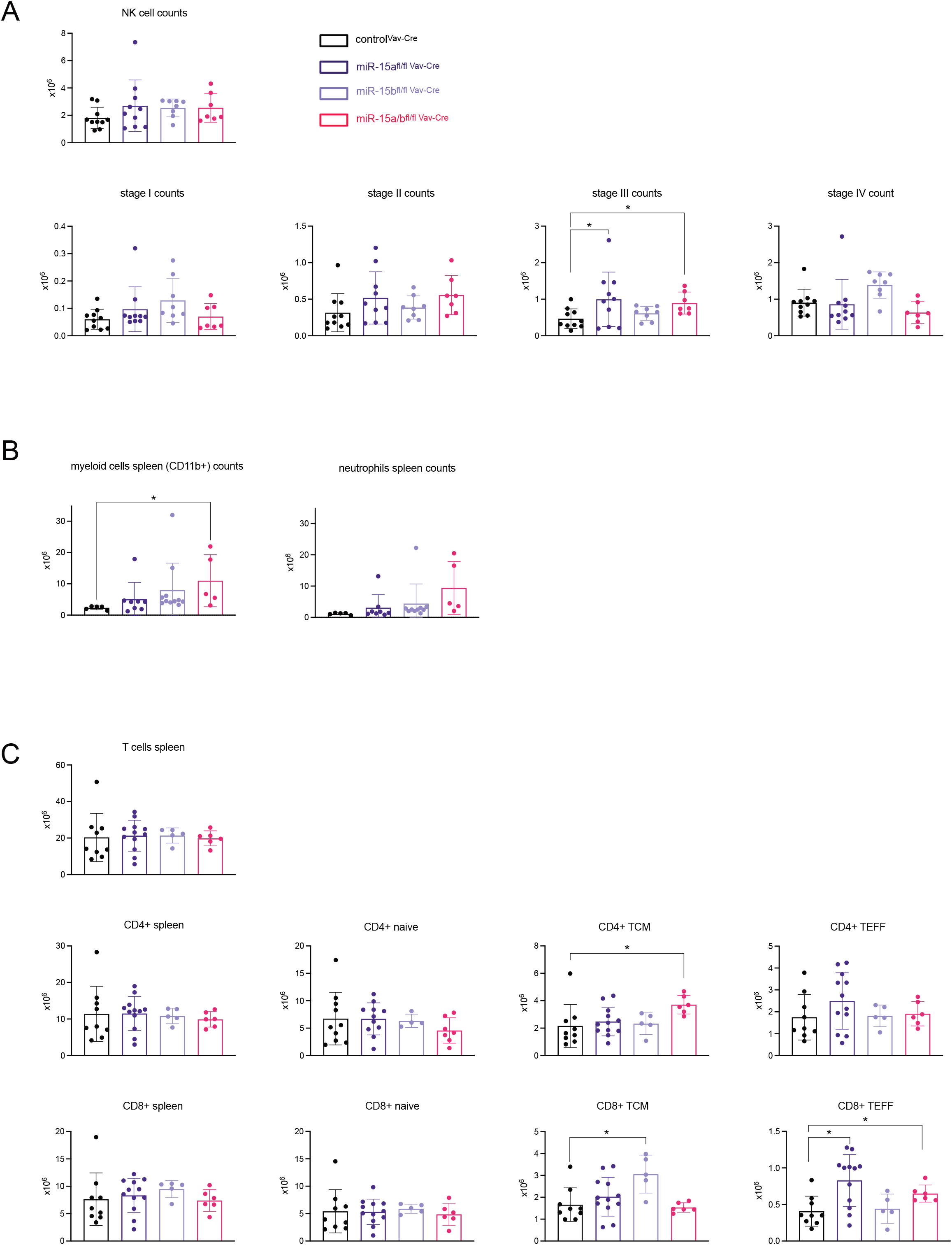
Total cell counts of splenic immune cell subsets. Total splenic cell numbers of indicated organs were counted and calculated accordingly. The bar graphs summarize the total cell counts for each described immune cell subset for miR-15a^fl/fl Vav-Cre^, miR-15b^fl/fl Vav-Cre^, miR-15a/b^fl/fl Vav-Cre^ and miR-15a/b^+/+ Vav-Cre^ mice. Each dot represents the data derived from one mouse. Error bars depict the standard deviation of the mean. Each of the different miR-15 knockout groups were statistically compared to miR-15a/b^+/+ Vav-Cre^ mice by an unpaired two-tailed Student’s t-test. *P < 0.05; *P < 0.005; ***P < 0.0005.

**Figure S2.**
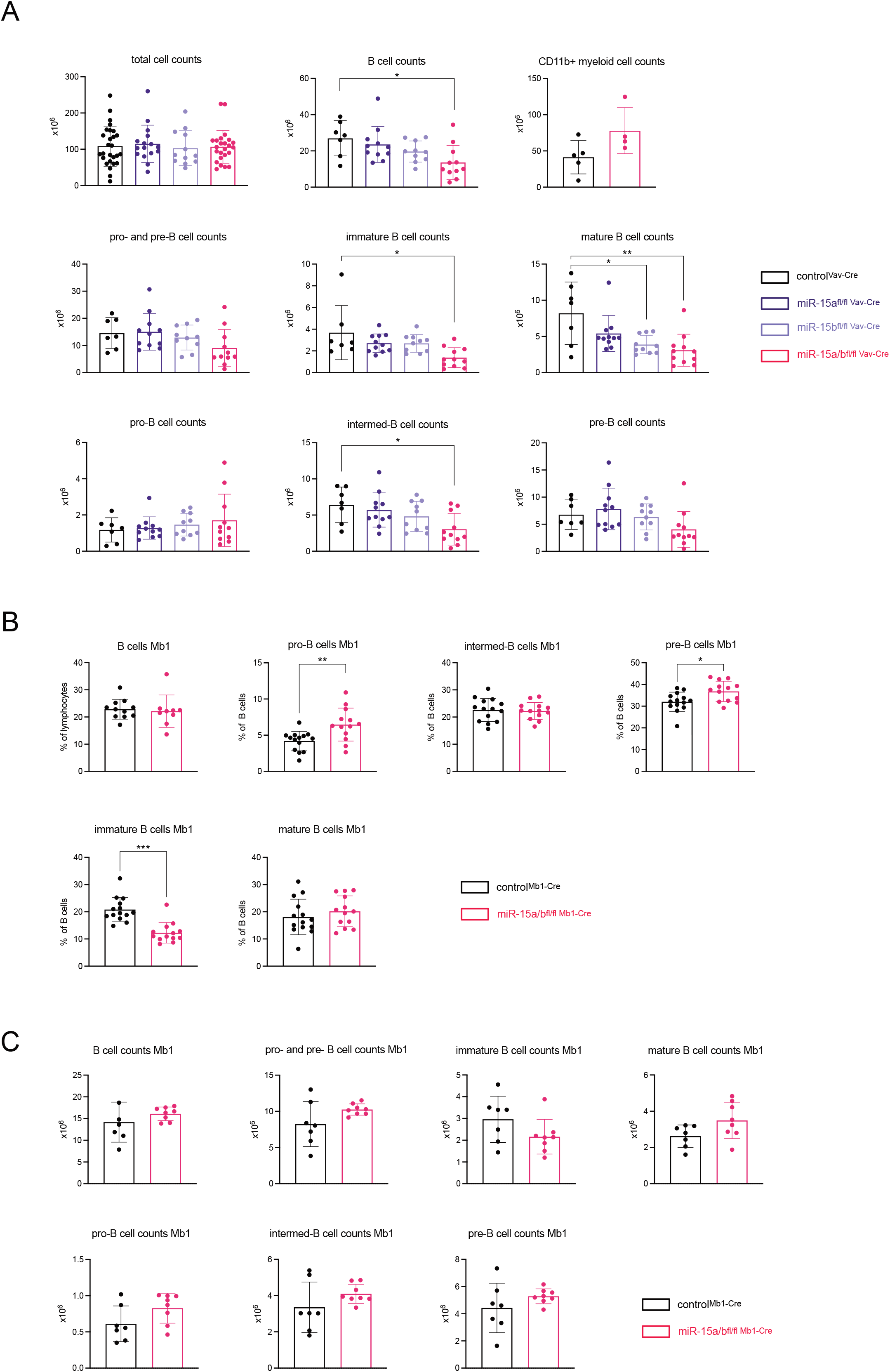
Total cell counts of bone marrow immune cell subsets. (A and C) Total bone marrow cell numbers of indicated immune cell subsets were counted and calculated accordingly. The bar graphs summarize the total cell counts for each described subset for miR-15a^fl/fl Vav-Cre^, miR-15b^fl/fl Vav-Cre^, miR-15a/b^fl/fl Vav-Cre^ and miR-15a/b^+/+ Vav-Cre^ mice (A) or miR-15a/b^fl/fl Mb1-Cre^ and miR-15a/b^+/+ Mb1-Cre^ mice respectively (C). (B) Bar graphs summarize the percentages of indicated bone marrow B cell subsets in miR-15a/b^fl/fl Mb1-Cre^ and miR-15a/b^+/+ Mb1-Cre^ mice. (A-C) Each dot represents the data derived from one mouse. Error bars depict the standard deviation of the mean. Each of the different miR-15 knockout groups were statistically compared to miR-15a/b^+/+ Vav-Cre^ mice by an unpaired two-tailed Student’s t-test. *P < 0.05; *P < 0.005; ***P < 0.0005.

**Figure S3.**
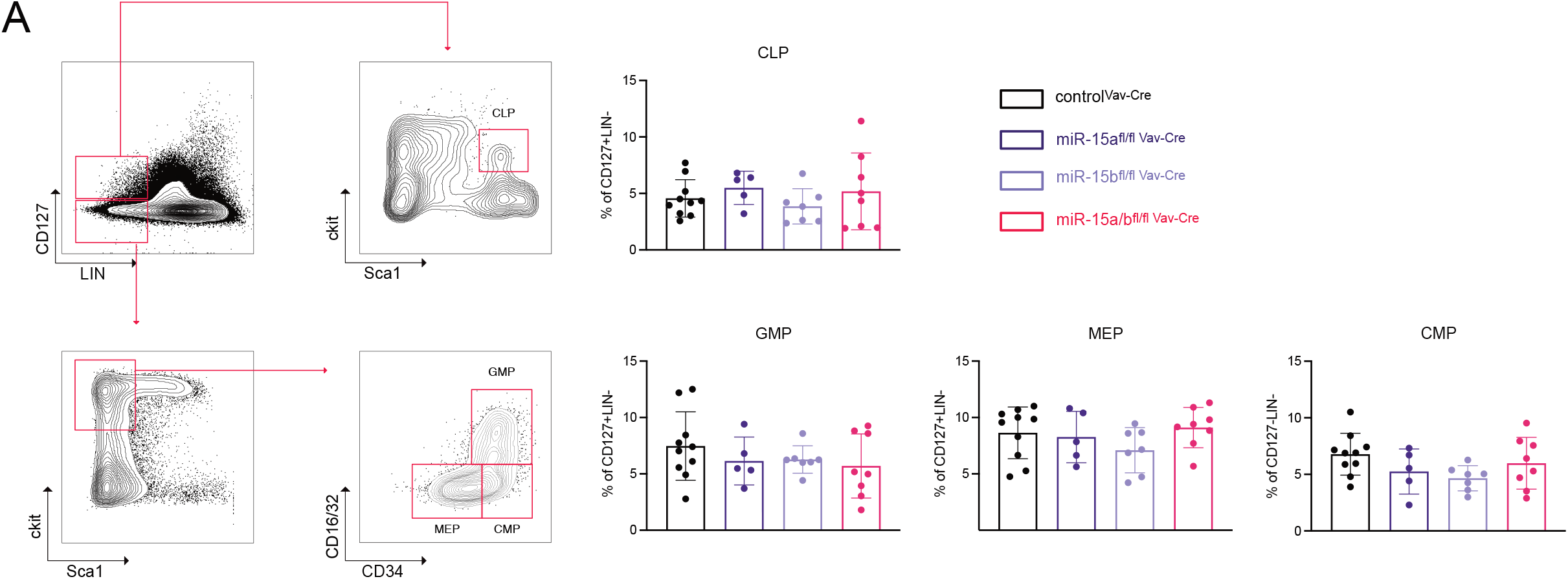
Flow cytometric analysis of CLPs, GMPs, MEPs, CMPs and B cell stages according to Hardy. (A) Schematic gating strategy for flow cytometric quantification of common lymphoid progenitors (CLPs), granulocyte-monocyte progenitors (GMPs), megakaryocyte-erythroid progenitors (MEPs) and common myeloid progenitors (CMPs) (left). The bar graphs summarize the percentages of indicated populations in the bone marrow of miR-15a^fl/fl Vav-Cre^, miR-15b^fl/fl Vav-Cre^, miR-15a/b^fl/fl Vav-Cre^ and miR-15a/b^+/+ Vav-Cre^ mice. Each dot represents the data derived from one mouse. Error bars depict the standard deviation of the mean. Each of the different miR-15 knockout groups were statistically compared to miR-15a/b^+/+ Vav-Cre^ mice by an unpaired two-tailed Student’s t-test. *P < 0.05; *P < 0.005; ***P < 0.0005.

**Figure S4.**
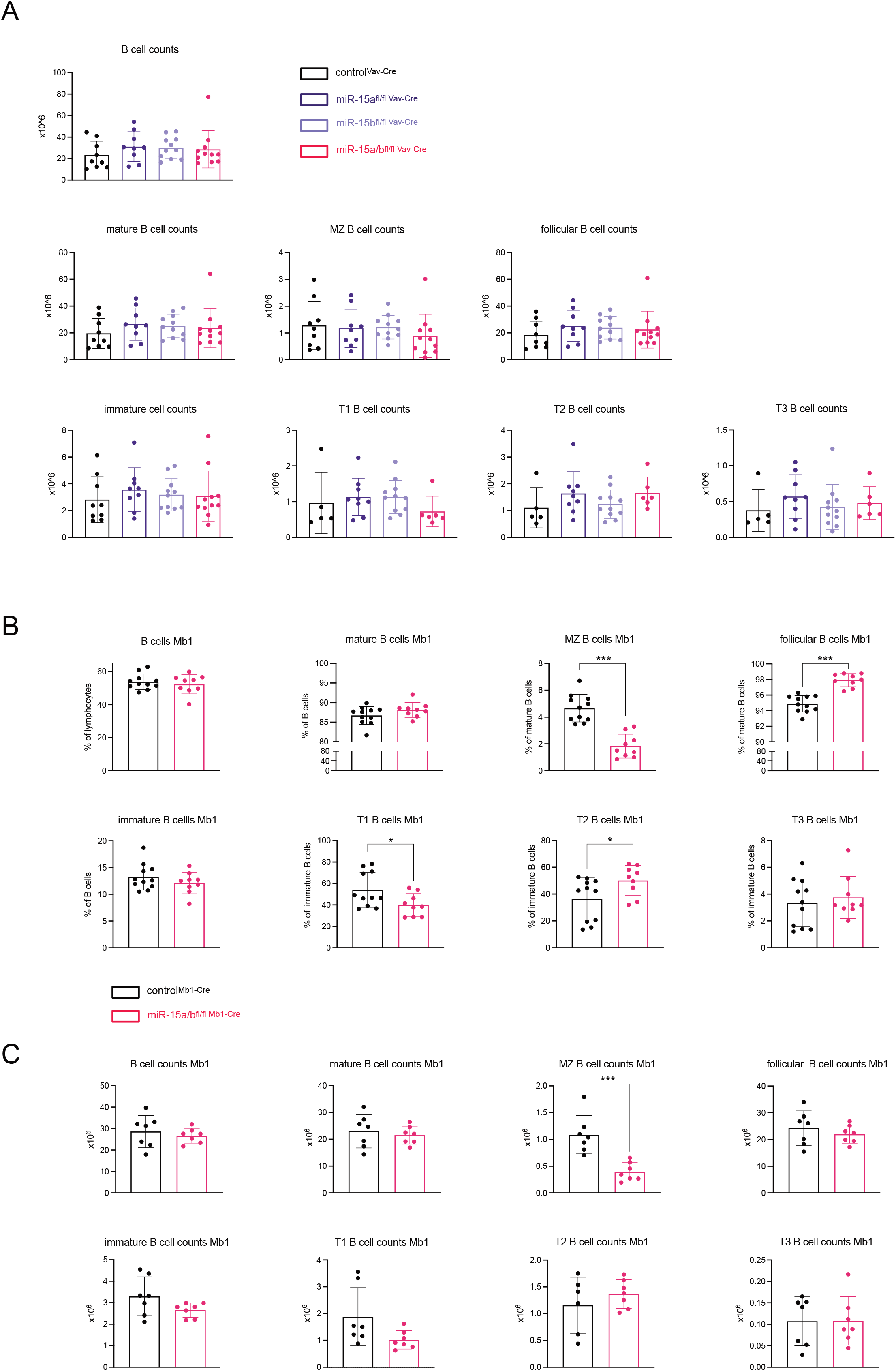
Splenic B cell numbers and analysis of splenic B cells in miR-15 knockout mice. (A and C) Total splenic cell numbers of indicated B cell subsets were counted and calculated accordingly. The bar graphs summarize the total cell counts for each described immune cell subset for miR-15a^fl/fl Vav-Cre^, miR-15b^fl/fl Vav-Cre^, miR-15a/b^fl/fl Vav-Cre^ and miR-15a/b^+/+ Vav-Cre^ mice (A) or miR-15a/b^fl/fl Mb1-Cre^ and miR-15a/b^+/+ Mb1-Cre^ mice respectively (C). (B) Bar graphs summarize the percentages of indicated splenic B cell subsets in miR-15a/b^fl/fl Mb1-Cre^ and miR-15a/b^+/+ Mb1-Cre^ mice. (A-C) Each dot represents the data derived from one mouse. Error bars depict the standard deviation of the mean. Each of the different miR-15 knockout groups were statistically compared to miR-15a/b^+/+ Vav-Cre^ mice by an unpaired two-tailed Student’s t-test. *P < 0.05; *P < 0.005; ***P < 0.0005.

**Figure S5.**
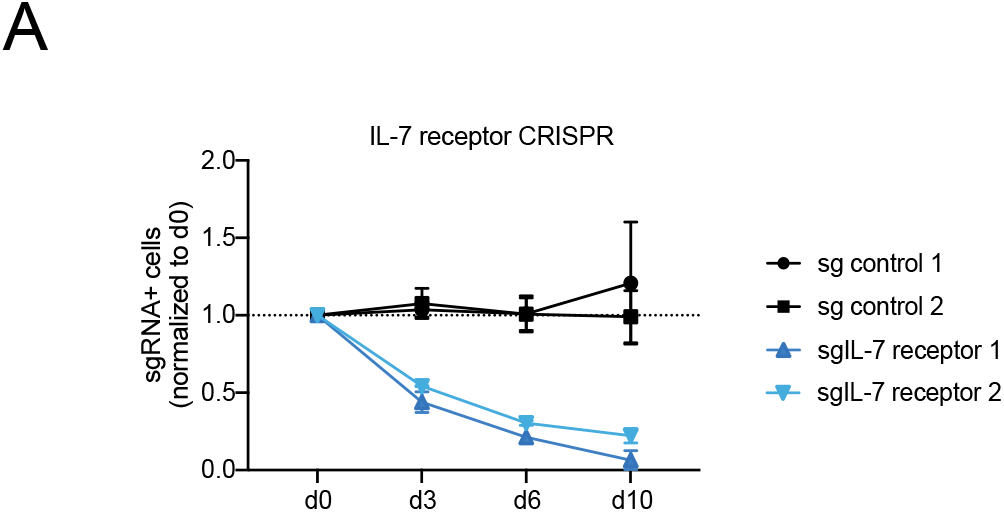
B cell progenitors depend on IL-7 receptor expression. Cas9 expressing progenitors were transduced with constructs coding for dsRed as a marker and guide RNAs targeting CD8 (sg control 1+2) or the IL-7 receptor (sg IL-7 receptor 1+2). The percentage of dsRed+ cells was analyzed for the respective time points by flow cytometric analysis and normalized to d0. The diagram depicts the mean percentages of dsRed+ cells for each day normalized to d0. Error bars depict the standard deviation of the mean.

